# The new phycobilisome linker protein ApcI regulates high light adaptation in *Synechocystis* sp. PCC 6803

**DOI:** 10.1101/2024.09.09.612062

**Authors:** Roberto Espinoza-Corral, Tomáš Zavřel, Markus Sutter, Chase H. Leslie, Kunwei Yang, Warren F. Beck, Jan Červený, Cheryl A. Kerfeld

## Abstract

Phycobilisomes are versatile cyanobacterial antenna complexes that harvest light energy to drive photosynthesis. These complexes can also adapt to various light conditions, dismantling under high light to prevent photo-oxidation and arranging in rows under low light to increase light harvesting efficiency. Light quality also influences phycobilisome structure and function, as observed under far-red light exposure. Here we describe a new, phycobilisome linker protein, ApcI (previously hypothetical protein sll1911), expressed specifically under red light. We characterized ApcI in *Synechocystis* sp. PCC 6803 using mutant strain analyses, phycobilisome binding experiments, and protein interaction studies. Mutation of *apcI* conferred high light tolerance to *Synechocystis* sp. PCC 6803 compared to wild type with reduced energy transfer from phycobilisomes to the photosystems. Binding experiments revealed that ApcI replaces the linker protein ApcG at the membrane-facing side of the phycobilisome core using a paralogous C-terminal domain. Additionally, the N-terminal extension of ApcI was found to interact with photosystem II. Our findings highlight the importance of phycobilisome remodeling for adaptation under different light conditions. The characterization of ApcI provides new insights into the mechanisms by which cyanobacteria optimize light-harvesting in response to varying light environments.

## Introduction

Harvesting of light energy in photosynthetic organisms is highly regulated to drive photosynthesis under diverse environmental conditions (Sanfilippo et al., 2019). In cyanobacteria, light is captured by water-soluble phycobilisomes (PBS), which are pigment-protein complexes that transfer the absorbed energy to the reaction centers of photosystems embedded in the thylakoid membrane (Gantt et al., 1968; Govindjee and Shevela, 2011). One example of antenna adaptation to specific light conditions is the formation of PBS rows under low light closely packed with photosystem II (PSII) increasing the efficiency of photochemistry (Ho et al., 2017; Rast et al., 2019). Likewise, light quality changes trigger adaptive mechanisms to optimize photosynthesis, such as state transitions, which consist in the regulation of energy allocation to PSII or photosystem I (PSI) to prevent saturation of the electron transport chain, leading to the production of reactive oxygen species (Tamary et al., 2012; Zavrel et al., 2024). Long-term acclimation processes in cyanobacteria include proteome changes, such as the reduction of the size of antenna complexes under intense light and the expression of the Orange Carotenoid Protein for the dissipation of excess of light energy captured by PBS (Kirilovsky and Kerfeld, 2016; Zavrel et al., 2019; Kerfeld and Sutter, 2024; Srivastava et al., 2024).

PBSs harvest light energy through bilins, pigments covalent bound to phycobiliproteins, which tune their spectroscopic properties (Beck, 2024) to ensure directionality of energy transfer from rods to the core and ultimately to the photosystems (Sil et al., 2022). The PBS from *Synechocystis* sp. PCC 6803 (hereafter referred as *Synechocystis*) consists of a tricylindrical core surrounded by 6 rods forming a hemidiscoidal arrangement composed by hexamers of phycocyanin (αβ)_6_ in the rods and allophycocyanin forming four toroid-shaped trimers (αβ)_3_ per cylinder (Adir et al., 2020; Dominguez-Martin et al., 2022; Bryant and Gisriel, 2024; Sauer et al., 2024). Linker proteins play an important role in the PBS structure by connecting the core cylinders together as well as by attaching the rods to the core (Bryant and Canniffe, 2018; Dominguez-Martin et al., 2022; Sauer et al., 2024).

PBS have been reported to undergo structural remodeling involving various proteins which tune the PBS light harvesting ability under different light quality changes, as well as playing a role in their interaction with photosystems (Bryant and Gisriel, 2024). The recently characterized PBS linker ApcG from *Synechocystis* has been shown to play a role in energy transfer from PBS to the photosystems (Espinoza-Corral et al., 2024). Indeed, this conserved linker has now also been identified in hemiellipsoidal PBS from red alga *Porphyridium purpureum* (also named Lpp2 in red algae) (You et al., 2023), confirming its interaction with PSII through its N-terminal extension. Furthermore, in *Synechocystis,* the PBS rods containing the linker protein CpcL are able to interact with PSI, which might regulate photosynthetic activity under stress conditions, such as iron deficiency (Watanabe et al., 2014; Shimizu et al., 2023; Zheng et al., 2023). Likewise, cyanobacteria express the chlorophyll binding protein IsiA under iron deficiency, forming rings around PSI (Guikema and Sherman, 1983; Burnap et al., 1993; Toporik et al., 2019). Additionally, low light conditions trigger the expression IsiX (homolog of IsiA) along with ApcD4 and ApcB3 which have been proposed to form antenna-like complexes interacting peripherally with PSI (Soulier et al., 2020; Soulier et al., 2022; Gisriel et al., 2023a). A fascinating adaptation of cyanobacterial photosynthesis is the expression of far-red light PBS, consisting in rodless bicylindrical cores exhibiting red shifted absorbance maxima as the specific proteins expressed under these conditions tune the spectral properties of the bilins (Gisriel, 2024; Gisriel et al., 2024).

Here we identify and characterize a new PBS linker. We discovered this protein, hypothetical protein sll1911 through sequence homology to the C-terminus, PBS-binding region of ApcG. We named this linker protein ApcI; ApcH was recently named in hepacylindrical PBS from *Anthocerotibacter* panamensis (Jiang et al., 2023). While the C-terminal domain of ApcI interacts with PBS, its N-terminal extension interacts with PSII, participating in energy transfer from PBS to PSII. Deletion of *apcI* in *Synechocystis* provides light tolerance at high light intensity compared to wild type (1000 μmol photons m^−2^·s^−1^). We propose that ApcI can substitute for ApcG in binding to the PBS and interacts with PSII under conditions that reduce of plastoquinone pool increasing the efficiency of photosynthesis. Our results describe a new linker protein and a new mode for the PBS to remodel its interaction with PSII under different environmental conditions.

## Results

### ApcI domain conservation among cyanobacteria

The structure of the PBS is defined by the association of phycobiliproteins and linker proteins, forming the main two PBS components namely the core and rods, which display in *Synechocystis* a hemidiscoidal shape (Adir et al., 2020; Dominguez-Martin et al., 2022; Bryant and Gisriel, 2024; Sauer et al., 2024). The interaction between linker proteins and phycobiliproteins is often found in conserved domains as it is for ApcE (Pfam00427 and Pfam00502) that defines the PBS-core type and CpcG1 (Pfam00427) mediating rods attachment to the core, among others (Ducret et al., 1994; Bryant and Canniffe, 2018; Dominguez-Martin et al., 2022). The recently described linker protein ApcG shows a conserved C-terminal PBS binding domain that interacts with ApcA at the membrane-facing side of the PBS core (Dominguez-Martin et al., 2022). Interestingly, when we searched cyanobacterial proteomes with only the PBS binding domain of ApcG, we were able to identify a large number of homologues (244 in 377 proteomes that contain ApcE, see **Supplementary Table S1** and Methods for details) that aligned well at the PBS binding domain but were very different from ApcG in their N-terminal part. In *Synechocystis* corresponded to the hypothetical protein Sll1911 which we named ApcI in accord with precedent in the literature (Jiang et al., 2023).

Analysis of the sequence conservation among the 244 ApcI homologues identifies three discrete regions in the protein: N-terminal domain, middle domain, and the PBS binding domain (**Figure 1B**). The C-terminal domain of ApcI resembles the ApcG PBS binding domain also from its structural prediction, adopting an alpha-helix conformation (**Figure 1C**). Other than the conserved C-terminal domain, ApcG and ApcI share no amino acid conservation.

**Figure 1.**
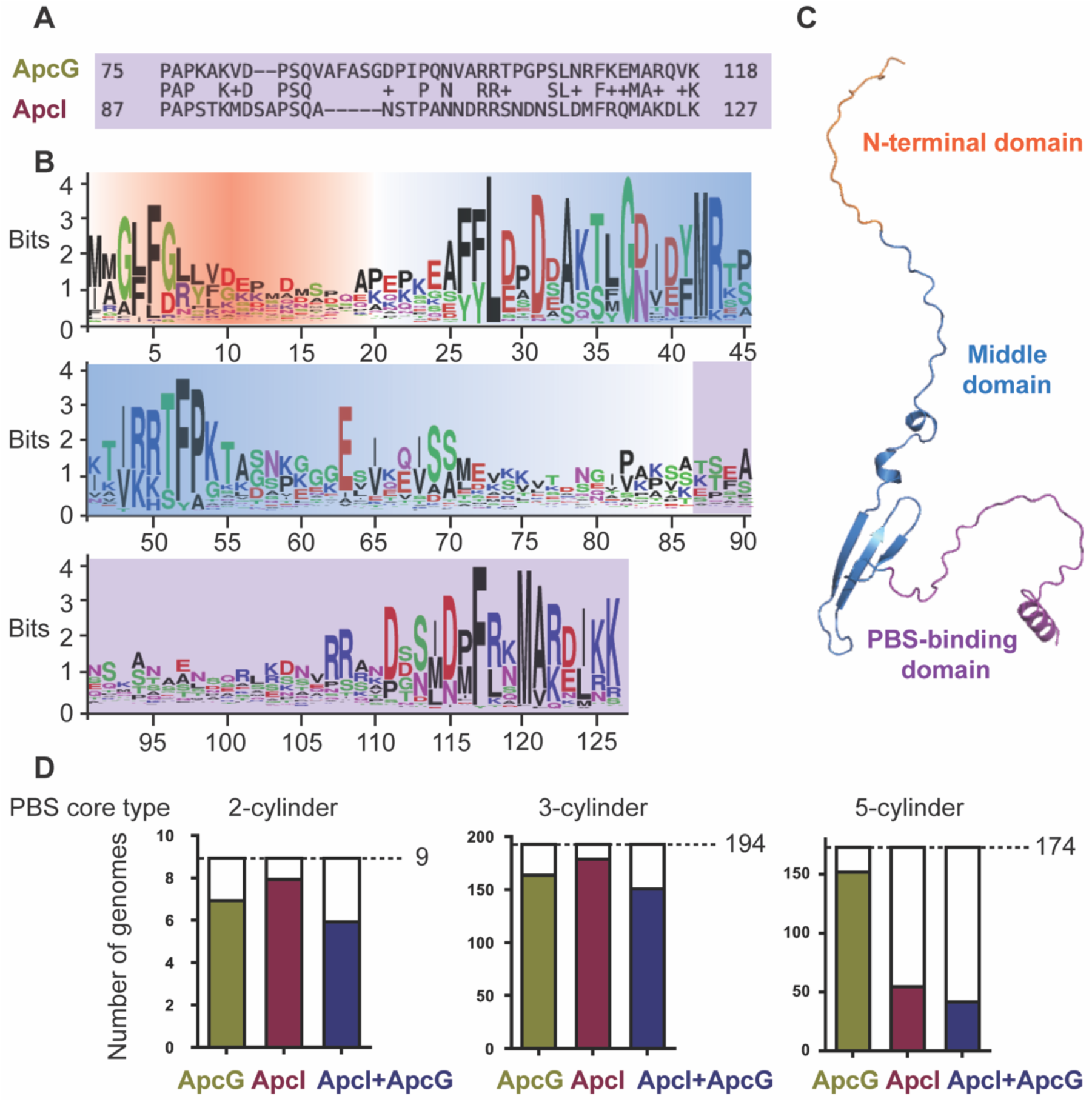
Sequence and structure overview of ApcI and its occurrence in cyanobacteria. (A) Sequence alignment of the C-terminal PBS-binding domains of ApcG and ApcI. (B) Amino acid sequence conservation logo based on 244 ApcI homologs. The N-terminal regionis highlighted in orange; the middle region is highlighted in blue, and the PBS-binding domain is highlighted in purple. (C) Alphafold structure prediction of ApcI highlighting the three regions, color coded as in (B). (D) A total of 377 ApcE-containing genomes categorized according to their PBS-core type using the length of *apcE* gene product as diagnostic. The number of genomes containing a gene encoding for either ApcG or ApcI is shown in bars as well as for the number of genomes with co-occurrence of genes encoding for both ApcG and ApcI.

However, as the ApcG N-terminal domains interact with photosystem II in the thylakoid membrane (Espinoza-Corral et al., 2024), we hypothesize that the N-terminal domain of ApcI could likewise interact with other complexes and tether the PBS to those. A structure Alphafold prediction (Jumper et al., 2021) suggests that the N-terminal domain of ApcI is unstructured while the middle domain, predicted to be beta strands could conceivably interact with a binding partner. The C-terminal PBS binding domain adopts an alpha-helix as expected from the similarity to the PBS-binding domain of ApcG (**Figure 1C**). Interestingly, the ApcG motif FxxM where the completely conserved phenylalanine interdigitates with ApcA is also conserved in the ApcI C-terminus (**Figure 1B**). These structural and sequence similarities between the ApcI and ApcG C-terminal domains strongly suggest the ability of ApcI to bind to the PBS core (Dominguez-Martin et al., 2022).

A sequence search of 377 cyanobacterial genomes that contain at least one gene encoding for ApcE shows that ApcI homologues are found in 64 % of the cyanobacterial genomes (244 out of 377) in contrast to ApcG which is found in 86 % of the genomes (325 out of 377) (**Supplementary table S1**). PBSs can be classified into different types by the length of ApcE (Bryant and Canniffe, 2018); they can either be bicylindrical (Glazer et al., 1979), tricylindrical (Bryant et al., 1979; Zheng et al., 2021) or pentacylindrical (Glauser et al., 1992; Ducret et al., 1998). Interestingly, ApcI is found more often in PBS with a bicylindrical and tricylindrical core (180 out of 194 and 8 out of 9 respectively) compared to pentacylindrical PBS (56 out of 174) (**Figure 1D**). In contrast, ApcG seems to be the more common PBS linker protein as it is present in the majority of the species analyzed regardless of their PBS core type (**Figure 1D**).

### Expression and physiological role of ApcI in light harvesting

To study the role of ApcI (*sll1911* gene locus) we generated a deletion strain replacing its native coding sequence with a chloramphenicol resistance cassette (Δ*apcI).* Furthermore, using the previously characterized Δ*apcG* strain (Espinoza-Corral et al., 2023), we generated a double mutant for both linker proteins replacing the native coding sequence of *apcI* by a kanamycin resistance cassette while having *apcG* gene replaced by a chloramphenicol resistance cassette (Δ*apcI,* Δ*apcG)* (**Supp. Figure S1**). Strains were grown under different white light intensities to compare their photo-inhibition at high light intensities. The Δ*apcI* strain showed phenotype similar to the wild type between 25 - 750 μmol photons m^−2^·s^−1^ and light tolerance at 1000 μmol photons m^−2^·s^−1^, while the Δ*apcG* strain showed an increased light tolerance compared to both wild type and Δ*apcI*. Strikingly, the double mutant strain (Δ*apcI*, Δ*apcG*) showed remarkable light tolerance in contrast to wild type, Δ*apcI* and Δ*apcG* strains for which specific growth rates started to decrease as a consequence of photo-inhibition from 750 to 1000 μmol photons m^−2^·s^−1^ (**Figure 2A**). Pulse amplitude modulation fluorometry measurements showed that the maximal electron transport rate (ETR_max_) decreased for wild type, Δ*apcI* and Δ*apcG* strains under high light while the double mutant maintained ETR_max_ constant throughout all light intensities (**Figure 2B**). Interestingly, the double mutant showed lower ETR_max_ compared to the other strains at low and middle light intensities (between 25 and 500 μmol photons m^−2^·s^−1^) suggesting impaired energy transfer from PBS to the photosystems as a consequence of the loss of both ApcG and ApcI (**Figure 2B**). Interestingly, non-photochemical quenching (qN, a parameter to diagnose light stress) measured in strains grown under 1000 μmol photons m^−2^·s^−1^ showed that wild type, Δ*apcI* and Δ*apcG* experienced similar levels of light stress unlike the double mutant strain where negligible qN was measured (**Figure 2C**). Whole-cell absorption spectra from all strains were similar when grown under normal conditions (i.e., 25 μmol photons m^−2^·s^−1^), however mutant strains showed increased absorbance between 400 to 500 nm (**Figure 2D**). To better resolve the spectral features, we recorded low temperature absorption spectra (77K), showing a similar trend that indicate carotenoid accumulation compared to wild type (**Figure 2E**).

**Figure 2.**
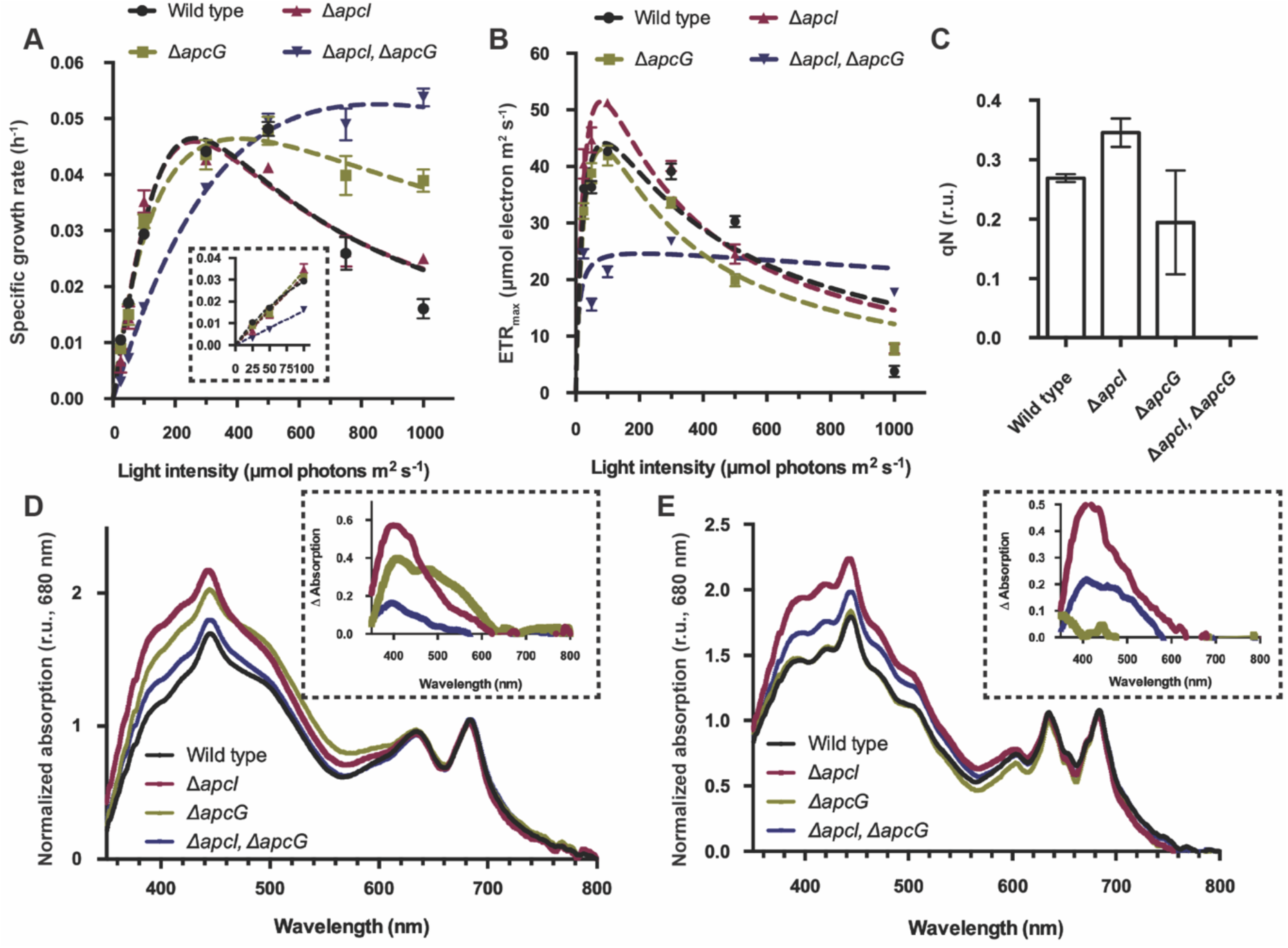
Physiological characterization of *apcI* strains. (**A**) Comparison of specific growth rate for Δ*apcG*, Δ*apcI* and double mutant strains with wild type under white light of intensities 25 - 1000 μmol photons m^−2^·s^−1^. Values correspond to averages of four biological replicates and error bars to SEM (standard error of mean). Fitting curves (Platt et al., 1980) are shown to model the behavior of the strains. Growth rates for the low light intensities are shown within a zoom-in dashed box. (**B**) Maximal electron transport rate (ETR_max_) comparison for single and double mutants with wild type. Values correspond to average of three biological replicates while error bars correspond to SEM. (**C**) Non-photochemical quenching (qN) from strains grown under 1000 μmol photons m^−2^·s^−1^. Values are presented as relative units (r.u.) (**D**) Whole-cell absorption spectra at room temperature from single, double mutants and wild type strains grown under 25 μmol photons m^−2^·s^−1^ of white light. Values correspond to normalized signals relative to their absorption at 680 nm. Curves represent averages of three biological replicates. (**E**) Whole-cell absorption spectra measured at 77K for wild type and mutant strains grown under 25 μmol photons m^−2^·s^−1^ of white light. Values correspond to averages of three biological replicates and presented as relative units (r.u.). Spectra in (**D**) and (**E**) are shown without error bars for clarity, and insets in (**D**) and (**E**) correspond to average spectra of mutant strains after subtracting the wild type absorption spectrum.

The high light tolerance exhibited by the Δ*apcI* strain compared to wild type (**Figure 2A**) suggests that ApcI could play a role in energy transfer from PBS to the photosystems at the thylakoid membrane. To test this, we compared low temperature fluorescence spectra (77K) from cells grown under white and red light, exciting chlorophyll (at 430 nm) and PBS (at 590 nm) separately. Chlorophyll emission spectra for cells grown under white light were indistinguishable among strains, indicating that PSII (680 nm peak) and PSI (720 nm peak) activities were unaffected by the lack of *apcI* (**Figure 3A**). Moreover, red light grown cultures (inducing state II) (Fuente et al., 2021) showed a higher PSII fluorescence for wild type and Δ*apcI* compared to Δ*apcG* and double mutant (Δ*apcI,* Δ*apcG*), consistent with the effect of ApcG in photosystem energy balance (Espinoza-Corral et al., 2024) (**Figure 3B**). Interestingly, when exciting PBS with white or red light grown cultures, PSII and PSI showed decreased fluorescence for all mutants compared to the wild type strain, indicating that the lack of either linker protein reduces the energy transfer from PBS to both photosystems (**Figure 3**). We tested if the light intensity difference between white (25 μmol photons m^−2^·s^−1^) and red (4 μmol photons m^−2^·s^−1^) lights could be the reason for the differences in chlorophyll emission spectra, however cultures grown under 4 μmol photons m^−2^·s^−1^ of white light showed similar spectra as for cells grown under 25 μmol photons m^−2^·s^−1^ (**Supp. Figure S2**). Additionally, we monitored the level of PSII, PSI and PBS protein accumulation by western blots using antibodies against their respective marker proteins. Interestingly, PSII and PBS levels were stable among strains grown under white or red light, while PSI showed reduction for wild type strain grown under red light compared to mutant strains (**Supp. Figure S3**). Therefore, we can conclude that the emission spectra differences among strains were not due to different levels of PBS or photosystems.

**Figure 3.**
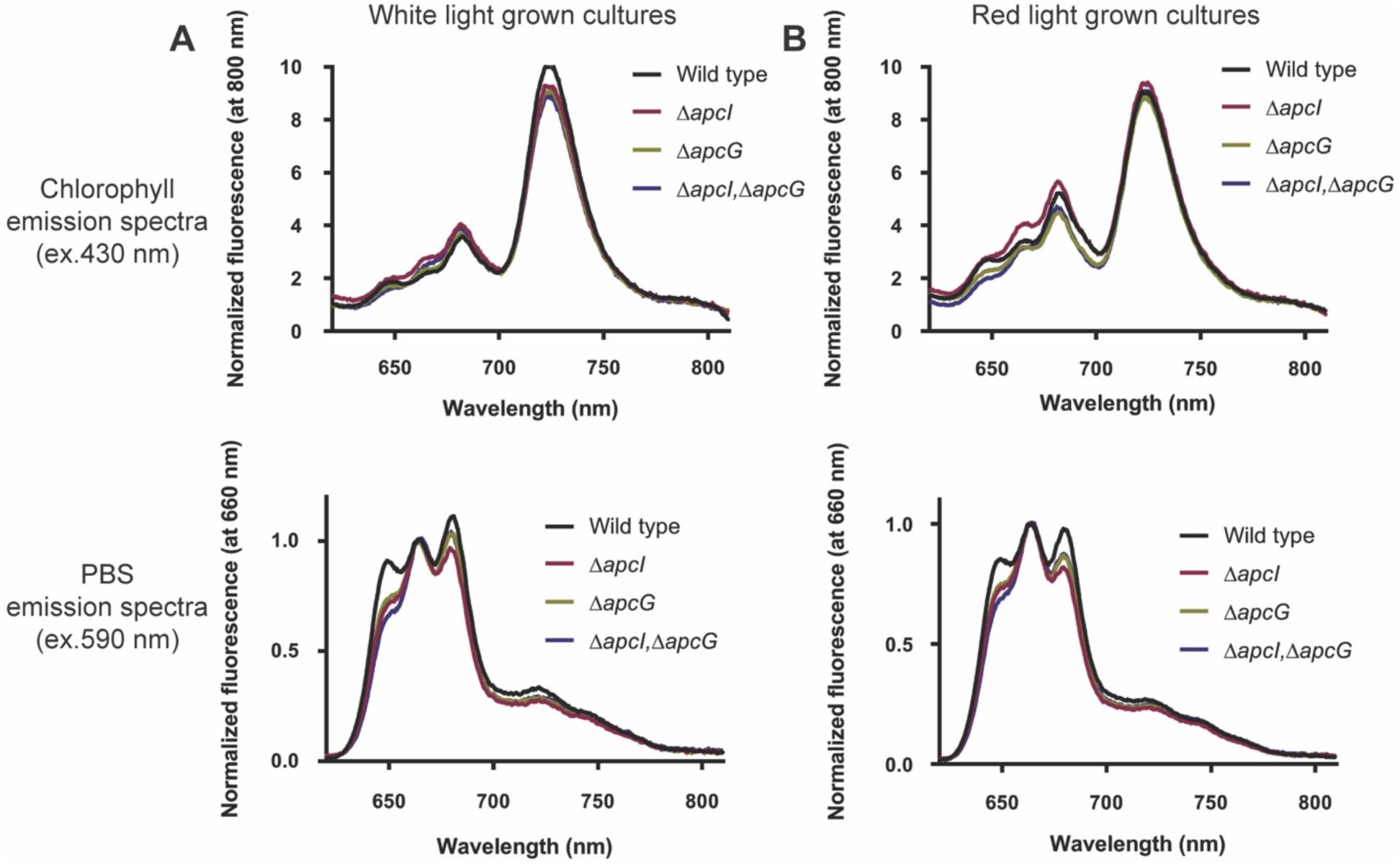
PBS energy transfer is impaired by the absence of ApcI. Cultures were grown under white light (25 μmol photons m^−2^·s^−1^) or red light (4 μmol photons m^−2^·s^−1^) for 4 days till they reached OD_720_ of 1–1.5 to record their whole-cell fluorescence emission spectra at 77K. Emission spectra of strains grown under white light are shown on panel (**A**), while those grown under red light are shown in (**B**). The spectra correspond to the mean of three biological replicates. Chlorophyll emission spectra were normalized by their fluorescence at 800 nm while PBS emission spectra were normalized using the PBS peak at 660 nm.

Proteomic analyses indicates that ApcI expression is induced under increasing light intensities (Zavrel et al., 2019). Additionally, transcriptomic data showed higher *apcI* transcripts in red light grown cultures (Luimstra et al., 2020). Using specific antibodies raised against full length ApcI (**Supp. Figure S4**), we identified that ApcI expression is indeed triggered by red light exposure (4 μmol photons m^−2^·s^−1^), while we could not detect ApcI in white light grown cultures (**Figure 4A**). Since red light exposure is known to trigger state II in cyanobacteria which is characterized by a highly reduced plastoquinone pool (Khorobrykh et al., 2020; Fuente et al., 2021; Zavrel et al., 2024), we hypothesized that the reduced plastoquinone pool in the thylakoid membrane is the trigger for ApcI expression. To test this, we cultivated wild type and Δ*apcI* strains under white light (25 μmol photons m^−2^·s^−1^) in the presence of DBMIB (an inhibitor of the cytochrome b_6_f complex preventing the oxidation of the plastoquinone pool) (Mao et al., 2002), thus chemically creating a potential trigger for ApcI expression. In the presence of DBMIB, we could detect ApcI from wild type cultures grown under white light, confirming that the plastoquinone pool redox state is likely the trigger for ApcI expression, rather than the specific light quality (**Figure 4B**). We then analyzed the localization of ApcI in *Synechocystis* cells by comparing membrane and soluble fractions from cultures grown under red light. Interestingly, ApcI was mainly found in the soluble fraction consistent with its putative PBS interaction while a small portion of it remained with the membrane fraction (**Figure 4C**).

**Figure 4.**
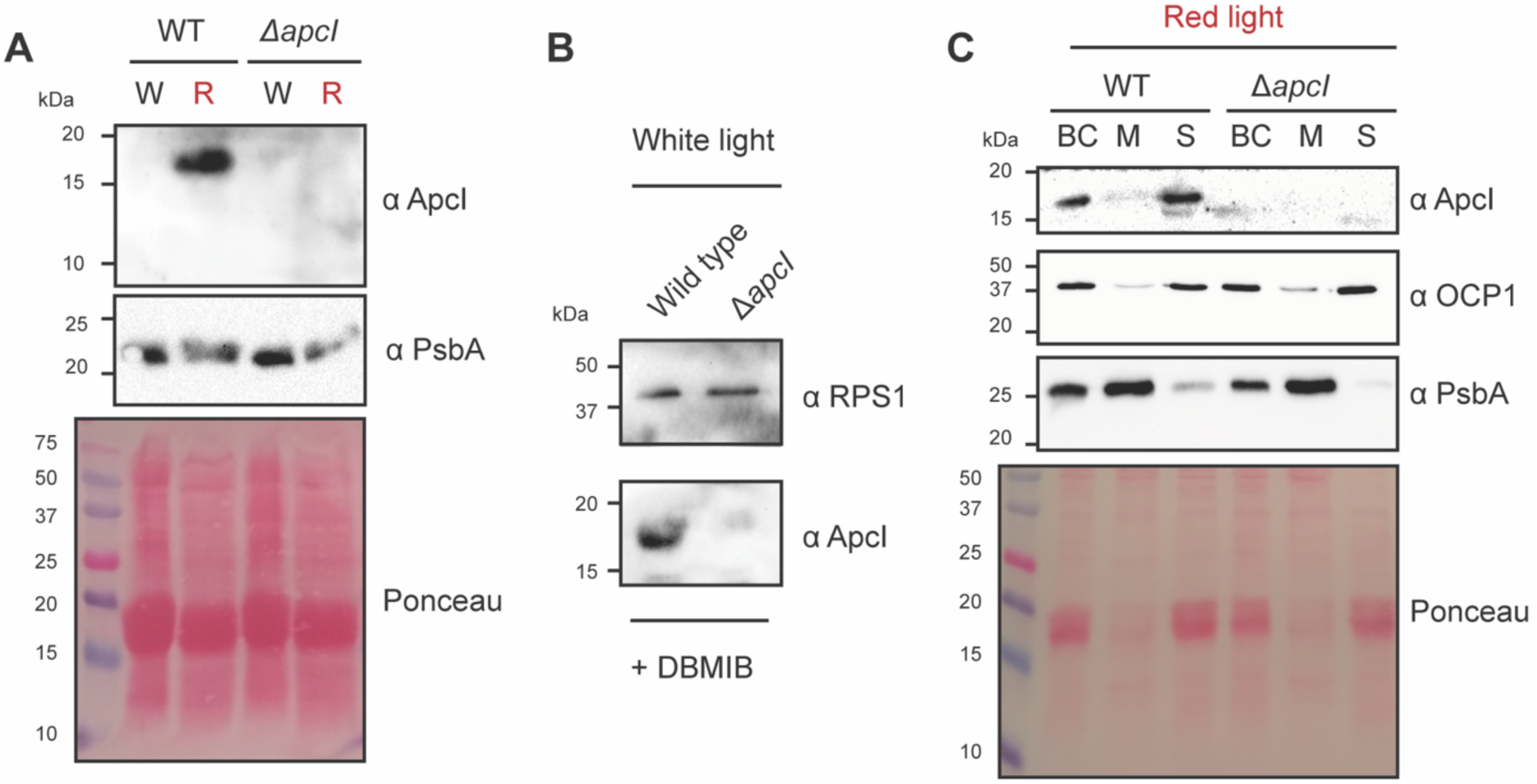
Expression of ApcI is induced by red light and a highly reduced plastoquinone pool. Expression of ApcI was monitored using antibodies raised against ApcI full length under different conditions. (**A**) Wild type and *apcI* deletion strains were grown under white light (W) or red light (R) to analyze the presence of AcpI in cell lysates. The subunit PsbA from PSII was used as a control. A total of 50 µg pf protein was loaded on each lane. (**B**) Wild type and *ApcI* deletion strains were grown under white light in the presence of DBMIB (50 µM) for 6 hours to detect ApcI in cell lysates. Antibodies against the protein RPS1 were used as protein loading control. A total of 25 µg of protein was loaded on each lane. (**C**) Strains were grown under red light for the expression of ApcI. Cell lysates (broken cells: BC) were further separated into membrane (M) and soluble fractions (S) by centrifugation. Antibodies against PsbA were used as a marker for the membrane fraction, while antibodies against OCP1 were used as a marker for the soluble fraction. Western blots shown correspond to a representative experiment out of three biological replicates.

### ApcI interacts with PBS and PSII

Because the C-terminal domain of ApcI is similar to the PBS binding domain from ApcG (e.g., FxxM motif which interdigitates with ApcA, **Figure 1A**) (Dominguez-Martin et al., 2022), we tested the possibility of ApcI binding to PBS *in vitro*. Isolated PBS from wild type and double mutant Δ*apcI*, Δ*apcG* were incubated with purified full length ApcI (**Supp. Figure S4A**) followed by discontinuous sucrose gradient centrifugation to separate excess unbound protein (located at the top of the sucrose gradient) and collect the intact PBS fraction (**Figure 5A**). Fractions containing PBS were precipitated by trichloroacetic acid (TCA) and analyzed by Western Blot using ApcI specific antibodies. Interestingly, ApcI was found to bind only to PBS in the double mutant strain Δ*apcI*, Δ*apcG* (**Figure 5B**), suggesting that ApcG bound to the wild type PBS prevents ApcI from interacting at the same site. To confirm that ApcI is binding to the pocket where ApcG binds to PBS using its C-terminal domain, we performed competition experiments using isolated PBS from double mutant strain Δ*apcI*, Δ*apcG* and the purified proteins ApcI and ApcG (**Supp. Figure S4**). When mutant PBS (Δ*apcI*, Δ*apcG*) were incubated with a mixture of ApcI and ApcG at equimolar ratio, ApcG prevented the binding of ApcI to PBS, supporting the hypothesis that these proteins bind to the same cavity at the PBS core (**Figure 5C**).

**Figure 5.**
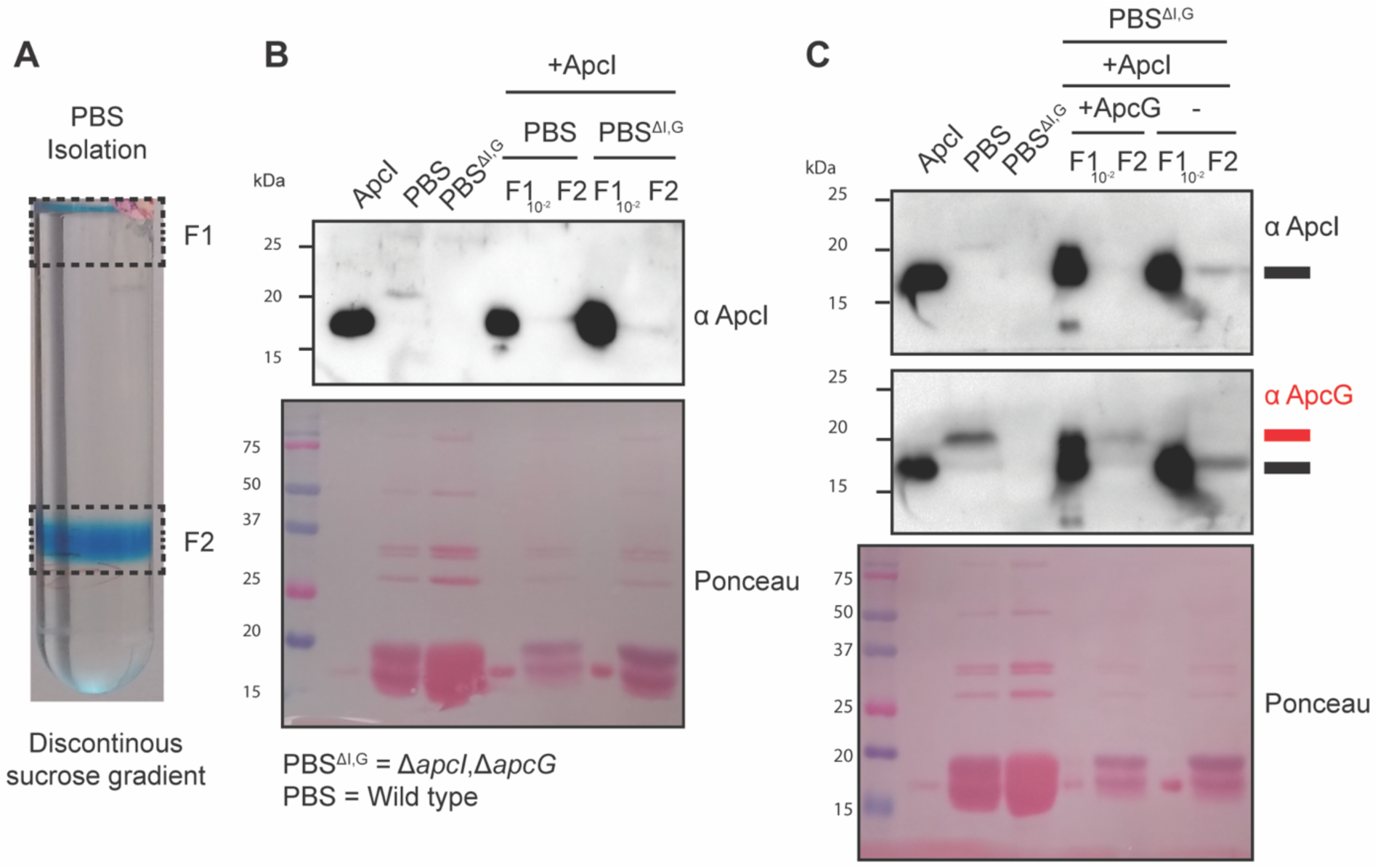
ApcI and ApcG interchangeable interaction with PBS core. Isolated PBS from wild type (denoted as PBS here) and double mutant (Δ*apcI*, Δ*apcG;* denoted as PBS^ΔI,G^) strains were used to perform binding assays with purified ApcG and ApcI. (A) Discontinuous sucrose gradient profile for the separation of intact PBS (fraction F2) discarding the excess of unbound protein (F1). (B) Wild type and mutant PBS were used in binding assays with purified ApcI. As a control, the purified protein ApcI was loaded along isolated PBS (precipitated by TCA) with fractions F1 (diluted to 10^-2^) and F2 (precipitated by TCA). (C) Competition binding assay using purified ApcG and ApcI. Fractions were loaded on the gel as described in (B). Nitrocellulose membrane was stripped after the detection of ApcI (highlighted in black) for its incubation with antibodies against ApcG (highlighted in red). Western blots shown correspond to a representative experiment out of three biological replicates.

While their C-terminal domains are structurally and functionally similar, the lack of similarity between the rest of the ApcI and ApcG suggests that these proteins play different roles, interacting with different partners. Low temperature fluorescence from Δ*apcI* showed impaired energy transfer to both PSI and PSII (**Figure 3**) suggesting it plays a role in the interaction between PBS and photosystems. To identify the interaction partner of ApcI at the thylakoid membrane, we designed a truncated version of ApcI, replacing its PBS binding domain with a His tag (ApcI^Δ74-128^-His) that allowed both purification of the protein as well as a method to pull down interaction partners (**Figure 6A**). However, incubation of ApcI^Δ74-128^-His with solubilized thylakoids (in the presence of 1 % dodecyl-beta-D-maltoside) induced aggregation of ApcI (**Supp. Figure S5A**). We then incubated ApcI^Δ74-128^-His with the soluble fraction from *Synechocystis* lysates and we found that the eluate from nickel beads turned green (**Supp. Figure S5B**). Western-blots analyses using the eluate from ApcI^Δ74-128^-His incubated with soluble fraction from cyanobacteria lysates showed the presence of the D1 protein of PSII (PsbA) but no detectable APC nor PSI proteins (PsaB) (**Figure 6B**). In order to confirm the interaction of ApcI with PSII, we generated a complementary strain for ApcI by replacing the native *psbA2* gene copy with *apcI* wild type open reading frame in the background of the Δ*apcI,* Δ*apcG* strain. This strategy ensured a strong expression of *apcI* under the control of the *psbA2* promoter (P*psbA2*) (Englund et al., 2016; Espinoza-Corral et al., 2024). Thylakoid complexes were separated in native gels to transfer the proteins to a membrane for the detection of ApcI using antibodies. Indeed, ApcI could be detected at the band where PSII is located (**Figure 6C**). To confirm this, we further separated the native gel bands by excising them from a native gel independently and further separating the proteins into a second SDS-PAGE dimension to detect ApcI by western-blots, showing that ApcI was found in the band were PSII is located (**Figure 6D**). This suggests that the N-terminal extension of ApcI binds to PSII in the thylakoid membrane.

**Figure 6.**
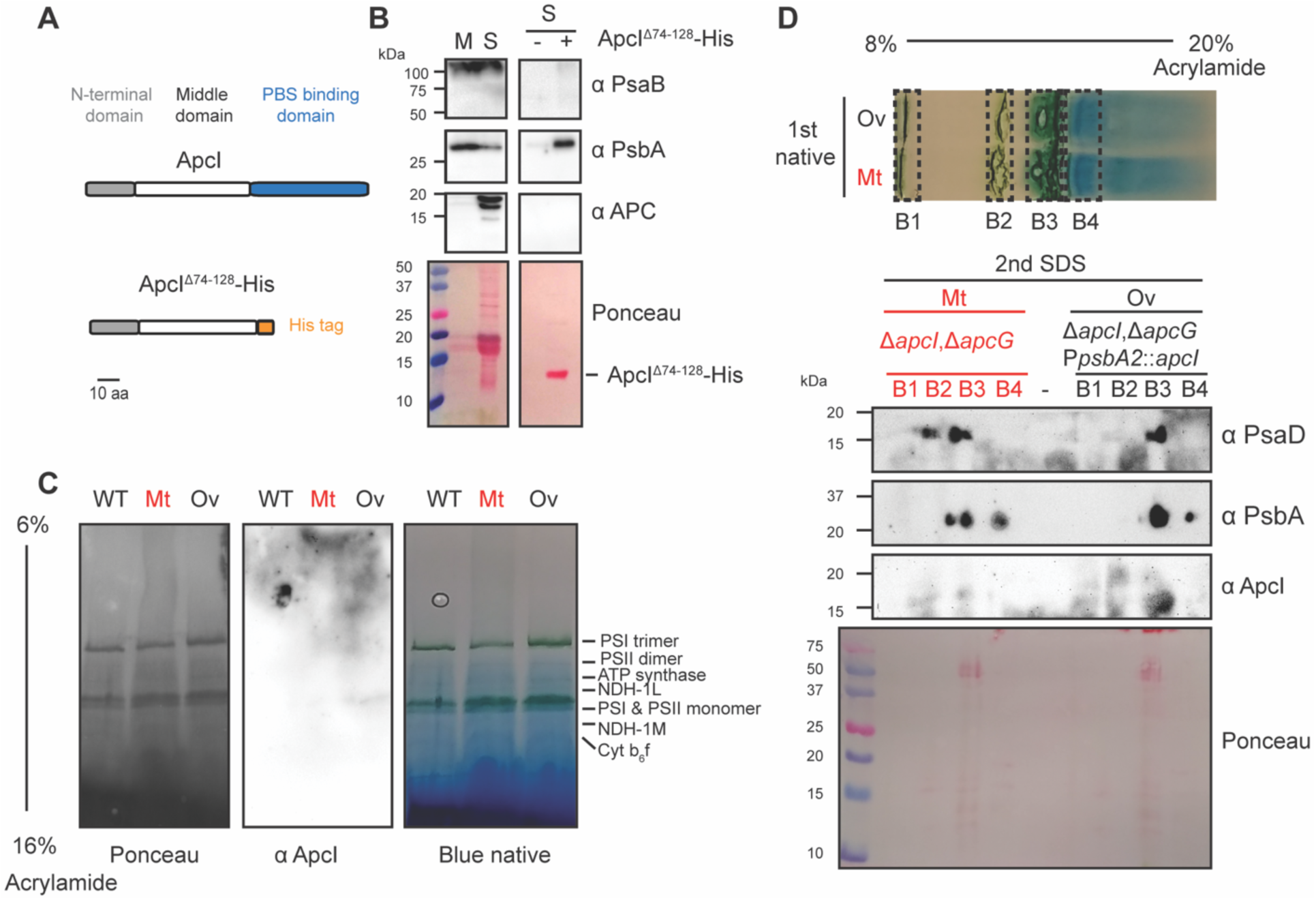
ApcI interacts with the PSII complex in the thylakoid membrane. (**A**) Truncated form of ApcI generated to perform pull-down experiments by replacing the PBS binding domain by a C-terminal His tag. (**B**) Pull-down experiments were performed preloaded with ApcI^Δ74-128^-His nickel beads and soluble protein from wild type *Synechocystis*. Antibodies against protein markers were used against PSI (PsaB), PSII (PsbA) and PBS (allophycocyanin; APC). As a control for antibodies, the membrane (M) and soluble (S) fractions were used from *Synechocystis* lysates. (**C**) The strains wild type, double mutant (Mt; Δ*apcI*, Δ*apcG*) and over-expressor (Ov; P*psbA2*::*apcI* in the background of Δ*apcI*, Δ*apcG*) were used to detect ApcI in blue native gels. (**D**) Blue native gels for the double mutant (Mt) and ApcI over-expressor (Ov) were performed to separate major protein complexes bands (B1, B2, B3 and B4) into a second denaturing dimension with SDS-PAGE. Western blots analyses used marker proteins for PSI (PsaD) and PSII (PsbA). Images correspond to a representative experiment out of three biological replicates.

## Discussion

Linker proteins play fundamental roles in regulating energy transfer between the PBS and the photosystems. Their distinct features may relate to the overall phycobilisome architecture as well as the environment in which they evolved. For example, the novel ApcH, acts as an anchor for the two extra core cylinders in heptacylindrical PBS from *Anthocerotibacter panamensis* (Jiang et al., 2023). Besides the terminal emitters ApcE and ApcD known to participate in energy transfer to PSII and PSI respectively (Gindt et al., 1992; Dong et al., 2009; Liu et al., 2013), recent cryogenic electron tomography structures of an algal PBS interacting with photosystems revealed additional linker proteins participating in the PBS-PS interaction (You et al., 2023). Under low light and far-red conditions, the cyanobacterial photosynthetic apparatus undergoes major remodeling along with PBS structure including the synthesis of chlorophyll *f* and *d* in PSII and rodless bicylindrical cores composed by specific phycobiliproteins that tune the absorption of bilins to drive photosynthesis under far-red light (Ho et al., 2016; Ho et al., 2017; Herrera-Salgado et al., 2018; Gisriel et al., 2023b). In *Synechocystis*, the linker protein ApcG participates in the PBS interaction with PSII as well as in regulating energy balance between photosystems (Espinoza-Corral et al., 2024), highlighting the role of linker proteins in PBS energy transfer. ApcI is the newest member of the family of PBS linker proteins and was discovered due to its PBS binding domain, which is similar to that of ApcG. The occurrence of ApcI among somewhat fewer cyanobacterial species compared to ApcG suggests that ApcI is a specialized, adaptive linker protein, while ApcG is the main PBS-PSII linker under normal conditions (**Supplementary table S1**). Indeed, *Synechocystis* proteomic data shows that under normal conditions (with a combination of red and blue photons and 25 μmol photons m^−2^·s^−1^ light intensity) ApcG is a hundredfold more abundant compared to ApcI (Zavrel et al., 2019). Additionally, increasing light intensity during *Synechocystis* cultivation reduces ApcG accumulation along with other antenna proteins while the expression of ApcI increases (Zavrel et al., 2019), suggesting a specific role for ApcI under high light conditions. Furthermore, ApcI occurrence in cyanobacterial species is often found in bi- and tri-cylindrincal PBS but less often in pentacylindrical (**Figure 1D**), suggesting there may be a correlation with PBS core structure.

Cyanobacterial growth under high light conditions (from 500 to 1000 μmol photons m^−2^·s^−1^ white light) is impaired as a consequence of photo-damage caused by the saturation of PSII (Nishiyama et al., 2006). However, comparison of wild type with single mutants Δ*apcI* and Δ*apcG* shows that the loss of function of either linker protein provides light tolerance under high light conditions with Δ*apcG* exhibiting higher light tolerance compared to Δ*apcI* (**Figure 2A**). Interestingly, *apcI* transcripts are more abundant under high light (Kopf et al., 2014), red light (Luimstra et al., 2020) and darkness (Saha et al., 2016) (conditions known to reduce the plastoquinone pool) (Fuente et al., 2021) compared to normal conditions (25 μmol photons m^−2^·s^−1^ white light), which suggests an adaptive role of ApcI under these conditions. Surprisingly, the double mutant Δ*apcI,* Δ*apcG* displayed remarkable light tolerance highlighting the role of these linker proteins in light harvesting (**Figure 2A**). Additionally, the Δ*apcI, ΔapcG* strain showed slower growth rate under lower light intensities compared to single mutants and wild type (from 25 to 300 μmol photons m^−2^·s^−1^ white light), consistent with the role of these linker proteins in the PBS-PS interaction (**Figure 2A**). Interestingly, the ETR_max_ of the double mutant Δ*apcI,* Δ*apcG* did not change under higher light intensities (**Figure 2B**) which could explain why this strain exhibits light tolerance as it might not be able to saturate the electron transport rate when missing both ApcG and ApcI linker proteins. Absorption spectra of strains grown under normal conditions (25 μmol photons m^−2^·s^−1^ white light) showed higher absorbance for mutant strains between 400 and 500 nm which was resolved by low temperature absorbance spectra measurements showing that the increase of absorbance (especially for Δ*apcI*) is very likely to be due to carotenoids accumulation (**Figure 2D-E**). Carotenoids are known to accumulate in cyanobacteria under stress conditions (Zakar et al., 2017; Rodrigues et al., 2023), which indicates that mutant strains Δ*apcI* and Δ*apcG* might experience stress due to the lack of the connectivity between PBS and photosystems.

The behavior of mutant strains compared to wild type suggests that their energy transfer from PBS to photosystems is reduced. In 77K emission spectra when exciting chlorophyll (at 430 nm), no differences were observed between wild type and Δ*apcI* strains grown under white or red light (**Figure 3**), in contrast to Δ*apcG* and double mutant Δ*apcI,* Δ*apcG,* which showed lower PSII emission when grown under red light triggering state II (Fuente et al., 2021), consistent with the role of ApcG in photosystem energy balance (Espinoza-Corral et al., 2024). On the other hand, PBS emission spectra (exciting at 590 nm) showed lower PSII and PSI fluorescence for all mutant strains compared to wild type under white or red light (**Figure 3**), indicating that ApcI is indeed necessary for the energy transfer from PBS to the photosystems. It is interesting that while Δ*apcG* showed differences in both chlorophyll and PBS emission spectra compared to wild type, Δ*apcI* only showed differences in PBS emission spectra, suggesting that only ApcG participates in regulating energy balance between photosystems. The expression of ApcI is shown to be triggered by a reduced plastoquinone pool, either induced by red light, or chemically under white light (**Figure 4**). Interestingly, comparison of growth and ETR_max_ under red cultivation light showed that the double mutant Δ*apcI,* Δ*apcG* did not show light tolerance as it happened under white cultivation light, while Δ*apcG* showed higher light tolerance compared to wild type and mutant strains (**Supp. Figure S6**). Under red light, Δ*apcI* performed slightly worse compared to wild type as well as to Δ*apcG*, which suggests a putative role of ApcI under light quality changes (**Supp. Figure S6**). Despite the more reduced PQ pool triggered by red light compared to white light illumination (Fuente et al., 2021), ApcI mediating PBS-PSII interaction does not lead to a PQ pool over reduction (**Supp. Figure S7**) suggesting no photo-damage under the conditions tested to trigger ApcI expression. On the contrary, it seems to optimize electron transport rate on the thylakoid membrane, which helps to keep ATP/NAPDH ratio stable over a wide range of light intensities (Hoper et al., 2024).

PBS binding and pulldown experiments showed that ApcI and ApcG bind to PBS in the same place (**Figure 5**), while the ApcI N-terminus (residues 1-73) binds to PSII (**Figure 6**). These results pose the question on the localization of ApcI, either attached to the PBS or at the thylakoid membrane. Localization of ApcI from strains grown under red light showed that the majority of ApcI is in the soluble fraction (**Figure 4C**). However, PBS isolated from strains grown under red light (that triggers the expression of ApcI, **Figure 4C**) did not have ApcI bound but ApcG (**Supp. Figure S8**). Nevertheless, the lack of ApcI still reduces the energy transfer from PBS to photosystems as shown in 77K emission spectra (**Figure 3**), which supports the ability of ApcI to bind to PBS and mediate its interaction with PSII (**Figure 5&6**). These observations strongly suggest that ApcI peripherally interacts with PSII at the thylakoid membrane, explaining its accumulation in the soluble fraction from cyanobacteria (**Figure 4C**) while absent in soluble PBS (**Supp. Figure S8**) and present in solubilized thylakoids (**Figure 6**). We summarize our observations as a model for the expression and interaction of ApcI with PSII where under normal conditions (balanced redox state of the plastoquinone pool) ApcG mediates PBS-PSII interaction, however under conditions that reduce the plastoquinone pool, such as red light, ApcG is replaced by ApcI (**Figure 7**). Interestingly, while both linker proteins do not share amino acid homology in their N-terminal and middle regions they both interact with PSII, suggesting that their interactions occur at different sites of the PSII dimer. Further investigation is required to identify the specific subunits from PSII that interact with either ApcG or ApcI.

**Figure 7.**
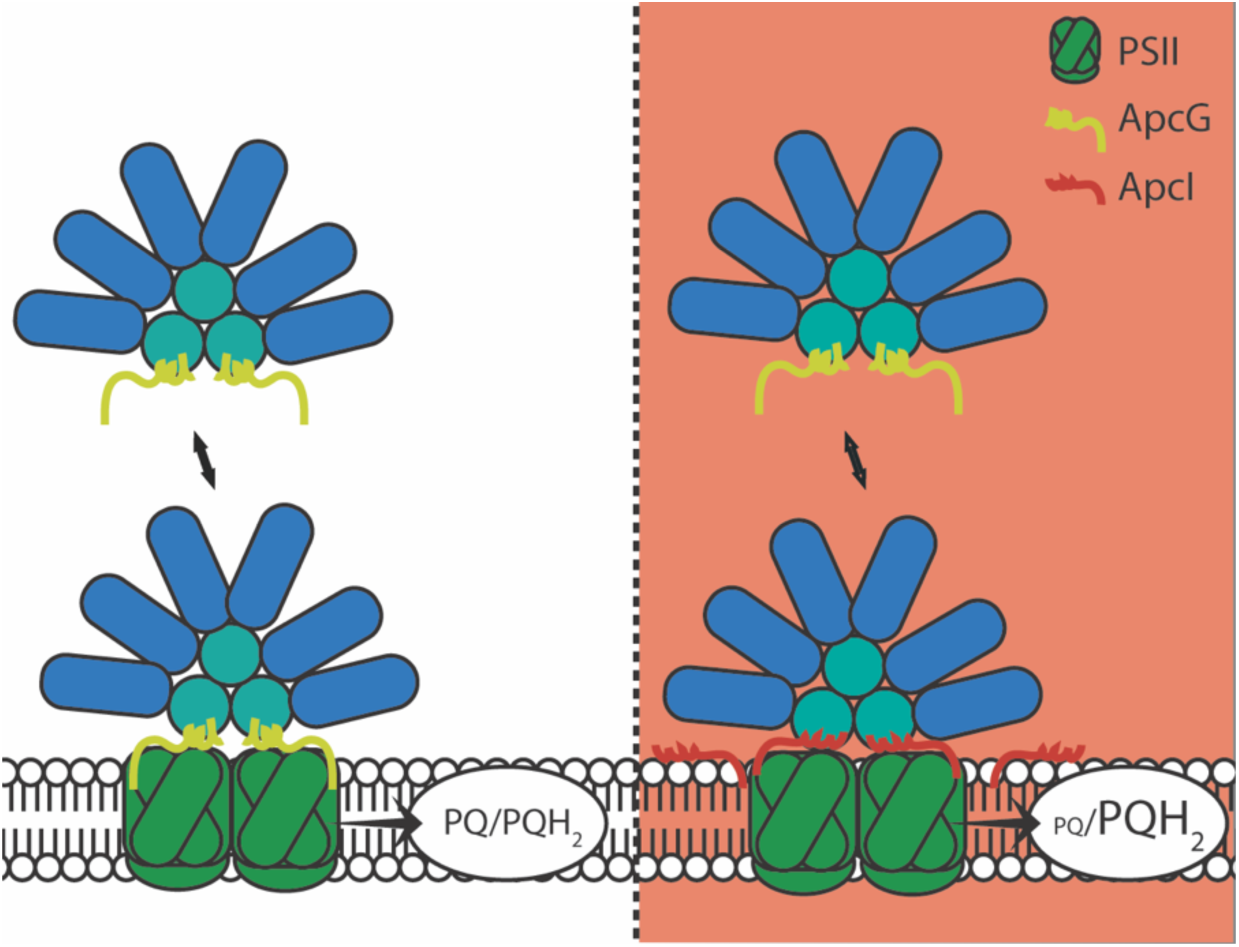
Model for ApcI expression and interaction with PSII. Under white light conditions (left panel), the plastoquinone pool is balanced between oxidized and reduced. However, under red light (right panel) the plastoquinone pool becomes more reduced, triggering the expression of ApcI (Figure 4) which remains associated with the thylakoid membrane via its N-terminal region (consisting of residues 1-73 which include N-terminal and middle domains) as evidenced using an ApcI over-expression strain (Figure 6). Upon PBS interacting with PSII, ApcG is exchanged by ApcI at the thylakoid membrane to interact with the bottom core of the tri-cylindrical through the C-terminal domain of ApcI (Figure 5).

## Materials and methods

### Bioinformatics and structure prediction

The sequences identified as ApcG in Dominguez-Martin et al. (2022) were truncated to their C-terminal PBS-binding domain (*Synechocystis* sp. PCC 6803, ApcG, *sll1873*, residues 75-121) and used to generate an HMM search model by aligning them with ClustalW (Thompson et al., 1994), trimming with trimAl (Capella-Gutiérrez et al., 2009) and the HMMs were generated using hmmbuild (http://hmmer.org/) (Potter et al., 2018). Then a curated list of cyanobacterial proteomes as described in Dominguez-Martin et al. (2022) containing a full-length ApcE (377 proteomes) was searched with hmmsearch (http://hmmer.org/) (Potter et al., 2018) to identify 244 protein homologues of ApcI (Sll1911 in *Synechocystis* sp. PCC 6803) (**Supplementary table S1)**. Cyanobacterial proteomes were further classified into bicylindrical, tricylindrical and pentacylindrical according to the length of their ApcE protein. The structural prediction for ApcI was obtained using AlphaFold prediction (Jumper et al., 2021).

### Cyanobacterial growth conditions

*Synechocystis* sp. PCC 6803 strains were cultivated in BG-11 medium (Rippka et al., 1979), buffered to pH 8 with 10 mM HEPES, at 28°C to 30°C, constant illumination (25-30 μmol photons m^−2^·s^−1^, white light) and supplemented with 3% CO_2_ (v/v) with agitation (160 rpm), corresponding to normal conditions. Cultures grown under red light (**Supp. Figure S9**) were cultivated in BG-11 medium as stated above but without CO_2_ supplementation. Selection of mutant strains was performed on BG-11 plates supplemented with 3 g/L sodium thiosulfate and solidified with 1.2% Difco agar (w/v). Antibiotics were supplemented to the selection media with chloramphenicol (25 μg/mL), kanamycin (50 μg/mL) or spectinomycin (20 μg/mL). In case of DBMIB (dibromothymoquinone) treatment, cyanobacteria cultures grown under normal conditions (OD_720_ 1-1.5) were supplemented with 50 µM of DBMIB and incubated under the same conditions for another 6 hours followed by protein extraction.

Growth curves were recorded in Multi-Cultivator MC-1000-MIX (Photon System Instruments, Czechia) in turbidostat regime (OD_720_ = 0.5 – 0.51, corresponding with ∼10^7^ cells mL^−1^) in BG-11 cultivation medium with ferric citrate as iron source, 50 µM EDTA (van Alphen et al., 2018) and with 17 mM HEPES buffer (pH ≈ 8). Temperature during all turbidostat cultivations was set to 30 °C. CO2 was supplemented by Gas Mixing System GMS-150 (Photon System Instruments) in final concentration 0.5 % (v/v), flow rate within each 80 mL cultivation tube was set to ∼ 40 mL min^-1^. Light was provided by orange (R615) and warm white (WW) LEDs of the Multi-Cultivator; intensities were set to 10 – 900 µE m^-2^ s^-1^. The cultures were kept under each particular condition for at least 24 h. This period was long enough to secure full metabolic acclimation (Rodrigues et al., 2023). Specific growth rates were calculated as described in Espinoza-Corral et al. (2024).

### Photosynthetic parameter measurements

After cyanobacteria growth stabilization in Multi-Cultivators, sampling was performed to measure rapid light curves in light-acclimated state (AquaPen, Photon System Instruments) from which maximum electron transport rate ETR_max_ and non-photochemical quenching coefficient q_N_ were derived according to the following equations:

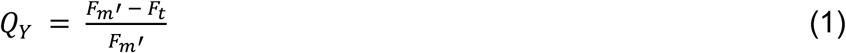

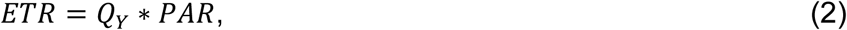

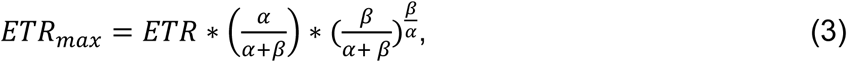

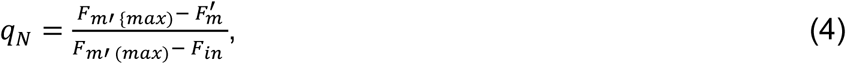

where F_m_’ and F_t_ are maximal and steady-state fluorescence in light-acclimated state, respectively, Q_Y_ is quantum yield of photosystem II (PSII; unitless), PAR is photosynthetically active radiation (units µmol photons m^-2^ s^-1^), ETR is electron transport rate (units electron m^-2^ s^-1^) (Ralph and Gademann, 2005), ETR_max_ is maximum electron transport rate, F_m’ max_ is maximal steady-state fluorescence throughout all tested light intensities of the rapid light curves, F_in_ is steady-state fluorescence at the onset of the rapid light curves measurement, and q_N_ is coefficient of non-photochemical quenching, related to flow of the captured light energy into heat. The coefficients α and β are slopes of the rapid light curves related to quantum efficiency of photosynthesis and photoinhibition, respectively, derived from light curves fitting (Platt et al., 1980).

In addition, fast fluorescence induction kinetic curves (OJIP) were measured. After sampling from Multi-Cultivators, the cultures were dark-acclimated for 15-20 min, and the OJIP kinetic was measured in MULTI-COLOR PAM (Walz, Germany) using 625 nm saturation pulse of intensity 2,000 µE m^-2^ s^-1^ and duration 600 ms. The parameter VJ, corresponding with redox state of PQ pool (Toth et al., 2007; Tsimilli-Michael et al., 2009) was calculated as:

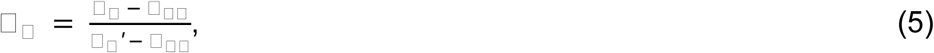

where FJ is fluorescence at the J-level of the OJIP curve, identified by the use of a second derivation of the fluorescence signal (Akinyemi et al., 2023) around 1 ms.

### Low-temperature fluorescence and absorption spectroscopy

Absorption spectra were recorded with a white probe beam and a multichannel CCD spectrometer in a fiber-optical spectrometer as described previously (Rose et al., 2023). Low temperature absorption spectra were recorded by decreasing the temperature stepwise to maintain cell integrity (**Supp. Figure S10**). Fluorescence spectra (**Supp. Figure S11**) were recorded with a home-built instrument (Gurchiek et al., 2020) employing a broadband LED and a compact double monochromator as an excitation source (2 nm spectral bandpass) and a detection system consisting of a 0.15 m spectrograph (4 nm spectral bandpass) and a back-illuminated CCD detector. Whole-cell samples suspended in a 60/40 (*v/v*) glycerol/BG-11 mixture (Espinoza-Corral et al., 2024) were held in 1 cm path length quartz cuvettes in a sample-in-gas liquid nitrogen cryostat (Oxford Instruments OptistatDN, with a MercuryiTC temperature controller). The absorption and fluorescence instruments were controlled by LabVIEW (National Instruments) programs. The spectra reported here are the average of those from three replicate samples.

### Generation and genotyping of *Synechocystis* strains

Wild type *Synechocystis* sp. PCC 6803 and Δ*apcG* strains (Espinoza-Corral et al., 2024) were used to generate *apcI* deletion strains. This was done by amplifying 500 bp upstream and downstream of *apcI* locus (*sll1911*) with primers oREC5 and oREC6 (**Supp. Table S2**) and cloning it into pJET1.2 generating plasmid pREC1. The *apcI* was deleted from pREC1 by inverse PCR using primers oREC7 and oREC8 introducing a *SacI* restriction site between the upstream and downstream regions of *apcI*, generating plasmid pREC2. A chloramphenicol resistance cassette as well as regions compatible for conjugation from pRL1075 (Black et al., 1993) were introduced into pREC2 using restriction site *SacI*, generating pREC3. Additionally, a second plasmid was generated for the incorporation of a kanamycin resistance cassette from pRL3313 into pREC2 using restriction site *SacI*, generating pREC8. A Bom site compatible for bacterial conjugation was incorporated into pREC8 using primers oREC48 and oREC49 generating pREC19. Over-expression of ApcI for complementation of mutant strains was obtained by cloning *apcI* gene using oREC14 and oREC15 into pET28a using restriction sites *XhoI* and *NdeI*, generating pREC5 which was used as template to amplify *apcI* with a T7 terminator region using primers oREC11 and oCK10, cloned into pPBSA2K5 (to drive the transcription of *apcI* using *psbA2* promoter, P*psbA2*) (Lagarde et al., 2000) using restriction sites *BamHI* and *NdeI* generating pREC7. A bom site compatible for conjugation was incorporated into pREC7 using primers oREC25 and oREC26 generating pREC10. Additionally, an *aadA* resistance cassette against spectinomycin was incorporated into pREC10 amplifying *aadA* from pRL3332 using primers oREC46 and oREC47 and restriction digestion using site *BamHI*, generating pREC18. Finally, a C-terminal His tag in *apcI* from pREC18 was removed using primers oREC50 and oREC51, generating pREC32. Wild type and Δ*apcG* strains were transformed by conjugation (Black et al., 1993) for the deletion of *apcI* gene using pREC3 (for generating single mutant Δ*apcI* with chloramphenicol resistance cassette) or pREC19 (transforming Δ*apcG* for the double mutant of Δ*apcG*, Δ*apcI* with chloramphenicol and kanamycin resistance cassettes). Genotyping of strains for *apcI* deletion was performed by extracting gDNA (Billi et al., 1998) and amplifying the wild type gene using oREC16 and oREC17, deletion with chloramphenicol cassette using primers oREC16 and oREC27 and deletion with kanamycin cassette using primers oREC16 and oREC44. Furthermore, the deletion of *apcG* was monitored amplifying *apcG* wild type using oREC12 and oREC13 and deletion with oREC12 and oREC27.

Likewise, complementation of the single mutant Δ*apcI* and double mutant Δ*apcG*, Δ*apcI* was performed by conjugation using plasmid pREC32 (with spectinomycin resistance cassette). Genotyping of *apcI* complementation was performed by amplifying a region containing *apcI*, *psbA2* promoter (P*psbA2*) and T7 terminator using primers oREC58 and oCK11 using as template gDNA from cyanobacteria.

### Protein expression and purification from *E. coli*

Expression of ApcI and ApcG was done by transforming BL21 DE3 (Invitrogen, Carlsbad, CA, USA) with the corresponding plasmids. For ApcI expression, the *apcI* gene from *Synechocystis* gDNA was amplified using primers oREC55 and oREC56 and cloned into a linearized pBbE2k vector using primers oCK23 and oCK24 by Gibson assembly (Gibson et al., 2009), generating pREC44 for the expression of ApcI with His-SUMO tag at its N-terminus under the control of tetracycline inducible promoter. In case of ApcG, the wild type *apcG* gene from *Synechocystis* gDNA was amplified with primers oREC59 and oREC60 (incorporating a His tag followed by a TEV site at the N-terminus of ApcG) and cloned by blunt ligation into pSL119 linearized with primers oCK3 and oCK4, resulting in pREC49. The recombinant ApcI^Δ74-128^-His protein started with sub-cloning *apcI* gene from pREC18 into pET11b using restriction digestion sites *BamHI* and *NdeI* which incorporated the His tag at the C-terminus of ApcI in pREC41. The truncation of ApcI deleting the PBS binding domain was performed by inverse PCR of pREC41 using primers oREC33 and oREC37, generating pREC45. Finally, the construct for His-[TEV]-ApcG expression was sub-cloned from pREC49 into pREC44 using restriction sites *NdeI* and *BamHI* generating pREC52 for the expression of His-[TEV]-ApcG under the control of tetracycline inducible promoter.

Expression of His-SUMO-ApcI was performed by transforming BL21 DE3 with pREC44 and using 4 liters of culture grown in luria broth under 37°C and induced when reaching OD_600_ ∼0.7 with 10 *μ*g/mL anhydrous tetracycline at 25°C overnight. Cells were centrifuged and resuspended in Buffer A (50 mM Tris pH 8, 200 mM NaCl) with protease inhibitor cocktail (Sigma, St. Louis, MO, USA), Dnase I (Sigma) and 50 mM imidazole followed by cell lysis using 2 passes through a cell disruptor (Constant Systems, Aberdeenshire, UK) at 15 kPSI. The soluble protein fraction was obtained by centrifuging the cell lysate for 30 min at 45,000 g and 4°C. The fusion protein His-SUMO-ApcI was purified by loading the cell lysate supernatant to a 5 mL HisTrap HP column (GE Healthcare, Little Chalfont, UK), washed with Buffer A, followed by a 5-column volume (CV) of 90% Buffer A and 10% Buffer B (v/v) (50 mm Tris pH 8, 200 mm NaCl, 500 mm imidazole) and eluted with a 5 CV gradient from 10% to 100% Buffer B (v/v). The purified His-SUMO-ApcI was incubated with ULP enzyme (purified in-home from plasmid pARH236 for the expression of fusion protein His-MBP-ULP) at a 1 to 20 ratio in Buffer A overnight at 4°C followed by a subtractive His trap purification using a 5 mL HisTrap HP column (GE Healthcare, Little Chalfont, UK) (GE Healthcare, Little Chalfont, UK) passing the mixture of His-SUMO-ApcI with ULP through the HisTrap column retaining His-SUMO as well as ULP and collecting the flowthrough containing tagless ApcI. Subsequently, ApcI was concentrated using a Amicon tube (Milipore) with 3 kDa cutoff and loaded onto a size exclusion column Superdex 200 increase 10/300 GL (Cytiva) using Buffer A at 4°C (**Supp. Figure S4A**). Finally, ApcI eluted from size exclusion chromatography at size of 32 kDa (**Supp. Figure S4B**) calculated using gel filtration standard (Bio-Rad, 1511901) (**Supp. Figure S4C**).

Expression of recombinant ApcI^Δ74-128^-His using pREC45 was done transforming BL21 DE3 and inducing 1 liter of cells with OD_600_ ∼0.7 and 1 mM IPTG overnight at 25°C. Cells were pelleted and resuspended in Buffer A (50 mM Tris pH 8, 200 mM NaCl) with protease inhibitor cocktail (Sigma, St. Louis, MO, USA), Dnase I (Sigma) and 50 mM imidazole followed by cell disruption using French press at 4°C. The soluble fraction was obtained by centrifugation for 30 min at 4°C and 45,000 x *g* and subjected to Histrap purification as described above. The elution from the Histrap column was further loaded onto an anion exchange resin (TOYOPEARL DEAE-650, CV 5 mL) using Buffer A with 20 mM NaCl collecting the flowthrough that contained the purified ApcI^Δ74-128^-His. Moreover, expression of the recombinant His-[TEV]-ApcG was done by transforming BL21 DE3 with pREC52 and using 1 liter of culture grown in LB at 37°C and induced when reaching OD_600_ ∼0.7 with 10 *μ*g/mL anhydrous tetracycline at 25°C overnight. Cells were pelleted and resuspended in Buffer A supplemented with protease inhibitor cocktail (Sigma, St. Louis, MO, USA), Dnase I (Sigma) and 50 mM imidazole followed by cell lysis using 2 passes through a cell disruptor. Cell lysate was centrifuged for 30 min and 45,000 x *g* at 4°C to obtain soluble protein fraction which was loaded onto a 5 mL HisTrap HP column. Elution of His-[TEV]-ApcG from Histrap column was performed as described above. Purified His-[TEV]-ApcG was incubated with TEV protease (purified in-home from plasmid pRK793, Addgene 8827) (Kapust et al., 2001) in Buffer A at a ratio of 1 to 20 and 4°C overnight. The digestion of His-[TEV]-ApcG with TEV protease was subsequently loaded onto a 5 mL HisTrap HP column collecting the flowthrough containing tagless ApcG (**Supp. Figure S4E**) following the same protocol above for ApcI. Finally, ApcG was further purified using cation exchange resin (TOYOPEARL SP-650, CV 5 mL) and performed the chromatography by gravity at 4°C as described in Espinoza-Corral et al. (Espinoza-Corral et al.). Protein concentration was measured using the BCA method (Pierce BCA Protein Assay Kit, 23227, Thermo Scientific).

### *Synechocystis* protein extraction

Cyanobacteria strains grown under normal conditions or red light were cultivated in 10 mL of BG-11 media in 25 mL flasks with agitation till they reached OD_720_ 0.5-1. Total protein extraction was obtained by centrifuging cyanobacteria cells followed by resuspension in extraction buffer at 4°C (50 mm HEPES pH 7.0, 25 mm CaCl_2_, 5 mm MgCl_2_, 10% [v/v] glycerol, and protease inhibitor cocktail). Subsequently, cells were broken by French press, followed by the addition of Triton X-100 1% (v/v) and incubation for 10 min on ice. Cell debris was discarded with 2 min centrifugation at 2,000 x *g* and 4°C. When separating soluble and membrane fractions from cyanobacteria cultures, cell pellets were resuspended in buffer TMK (50 mM Tris pH 7.5, 10 mM MgCl_2_ and 10 mM KCl) with protease inhibitor cocktail (Sigma, St. Louis, MO, USA). After cell disruption by French press, intact cells were discarded in the pellet by centrifuging the samples for 1 min at 2,000 x *g* and 4°C and the supernatant was further centrifuged for 30 min at 20,000 x *g* and 4°C. Supernatant was rescued corresponding to the total soluble protein fraction and pellet was resuspended in buffer TMK corresponding to membrane fraction. Protein concentration was measured using the BCA method (Pierce BCA Protein Assay Kit, 23227, Thermo Scientific).

### Immunoblot analysis

Proteins were separated into SDS-PAGE gels and transferred to a nitrocellulose membrane (Amersham, Protran) followed by blocking with 5% fat-free milk (w/v) in TBS (Tris 20 mm and 150 mm NaCl) at room temperature for 1 hour. Incubation of primary antibodies was done overnight at 4°C in TBS-T (TBS with 0.01% tween-20 [v/v]) (anti-PsbA; AS05 084A, anti-PsaB; AS10 695, anti-APC; AS08 277, anti-RPS; AS08 309, Agrisera). Membranes were washed 3 times with TBS-T for 15 min at room temperature and incubated with secondary polyclonal anti-rabbit antisera HRP for one hour at room temperature (Jackson ImmunoResearch, 111-035-003), followed by 3 additional washes with TBS-T and visualized by enhanced chemiluminescence technique.

Primary antibodies against ApcG and ApcI were raised immunizing rabbits with the purified proteins (**Supp. Figure S4**) (Genscript). Bleedings from rabbits were used in dilutions of 0.25 mL into 1 mL of TBS-T. Antibodies against OCP were used as described in Wilson et al. (2012).

### Pull-down experiments

Wild type *Synechocystis* cultures of 1 liter grown in BG-11 for 1 week under normal conditions were centrifuged and resuspended in 0.1 M phosphate buffer and pH 7.5 with protease inhibitor cocktail and 50 mM imidazole (Sigma, St. Louis, MO, USA) at 4°C. Cells were broken by French press with 4 passes followed by a first centrifugation of 1 min and 2,000 x *g* to discard intact cells and a second of 30 min and 45,000 x *g* at 4°C. The soluble fraction was rescued, and the membrane fraction was solubilized with 10 mL solubilization buffer (1% dodecyl-beta-d-maltoside [w/v], 750 mm aminocaproic acid, 50 mm Bis-Tris pH 7-, and 50 mM imidazole) followed by an incubation of 30 min on ice. Solubilized membranes (majority thylakoids) were centrifuged for 30 min at 30,000 x *g* and 4°C discarding the pellet (insoluble complexes).

Pull-down experiments were performed using the purified ApcI^Δ74-128^-His pre incubated in NTA nickel beads (0.8 mL CV) with either soluble proteins or solubilized thylakoids. Beads were incubated for 1 hour at 4°C and gentle rotation. Non-interacting proteins were washed off from the beads by centrifuging them for 2 min and 100 x *g* at 4°C followed by 4 washes using 10 CV of either 0.1 M phosphate buffer and pH 7.5 with 50 mM imidazole (when using soluble protein fraction) or solubilization buffer (when using solubilized thylakoids). Elution was obtained by washing the beads with 0.1 M phosphate buffer and pH 7.5 with 200 mM imidazole or solubilized buffer with 200 mM imidazole.

### Separation of protein in first native dimension and second denaturing dimension gels

Samples from pull down experiments using either soluble proteins or solubilized thylakoids as well as solubilized thylakoids from different strains were loaded onto gradient native gels as described by Schagger and Vonjagow (1991). Native gels were run without Coomassie brilliant blue as described by Espinoza-Corral et al. (2024). Second denaturing dimension was done separating the proteins from native gels into 12% SDS-PAGE gels supplemented with 4 M urea. Proteins were visualized using the method described by Blum et al. (1987).

### PBS binding assays

Cyanobacteria PBS from wild type and mutant strains were isolated following the method described in Espinoza-Corral et al. (2024). Isolated PBS were incubated for 4 hours and 15°C under gentle rotation with either purified ApcI or a mixture of ApcG and ApcI (equimolar ratio) at a molar concentration of PBS/linker of 0.0015 using 15 pMol of PBS and 10000 pMol linker protein in 0.8 M phosphate buffer (pH 7.5). Samples were loaded onto discontinuous sucrose gradients of 1.5, 1, 0.75, 0.5, and 0.25 m phases in phosphate buffer (0.8 M, pH 7.5) and centrifuged overnight at 22°C and 25,000 rpm to separate unbound proteins (top of the gradient) from PBS bound proteins (between 0.75 M and 1 M sucrose phases). Fractions were rescued from the sucrose gradients (F1 unbound proteins at the top and F2 corresponding to PBS bound proteins) and precipitated by TCA for further analyses.

### Accession numbers

Sequence data from this article can be found in **Supp. Table S1** containing proteins IDs for Uniprot library.

### Software

Figures were produced using Adobe Illustrator CS6 and GraphPad Prism version 6.0 (GraphPad Software, La Jolla, CA, USA; www.graphpad.com). Semiquantitative analyses of western blots were generated using ImageJ (https://imagej.nih.gov). The sequence conservation logo was generated with Weblogo (<u>Crooks et al. 2004</u>) using trimmed sequences by trimAl (Capella-Gutiérrez et al., 2009). AlphaFold2 was used for structure prediction of ApcI (Jumper et al., 2021). Analysis of the rapid light curves and OJIP curves fluorescence data was performed with the use of own-built Python scripts; the tool is available online at https://tools-py.e-cyanobacterium.org.

## Author contributions

R.E.-C. designed and conducted the research, analyzed the data, and wrote the article; T.Z. conducted cyanobacteria growth experiments and photosynthetic parameter measurements and analyzed the data; J.C. analyzed the data; C.L and K.Y conducted low-temperature absorption and fluorescence spectra measurements and W.F.B. analyzed the data; C.A.K. and M.S. designed research, analyzed the data, and wrote the article; all authors provided comments on the manuscript and contributed to experimental design.

## Acknowledgments

Research in the Kerfeld lab was supported by the Office of Science of the U.S. Department of Energy under award number DE-SC0020606. Work in the laboratory of W.F.B. was supported by grant award DE-SC0010847 from the Photosynthetic Systems program of the Office of Basic Energy Sciences, U.S. Department of Energy.

## Abbreviations

PS: photosystem

PBS: phycobilisomes

APC: allophycocyanin

